# Statistical model integrating interactions into genotype-phenotype association mapping: an application to reveal 3D-genetic basis underlying Autism

**DOI:** 10.1101/2020.07.27.222364

**Authors:** Qing Li, Chen Cao, Deshan Perera, Jingni He, Xingyu Chen, Feeha Azeem, Aaron Howe, Billie Au, Jun Yan, Quan Long

**Affiliations:** Department of Biochemistry and Molecular Biology, University of Calgary, Alberta, Canada; Heritage Youth Researcher Summer Program, University of Calgary, Alberta, Canada; Department of Physiology and Pharmacology, University of Calgary, Alberta, Canada; Department of Medical Genetics, University of Calgary, Alberta, Canada; Department of Mathematics and Statistics, University of Calgary, Alberta, Canada; Alberta Children’s Hospital Research Institute, University of Calgary, Alberta, Canada; Hotchkiss Brain Institute, University of Calgary, Alberta, Canada

**Keywords:** Genotype-phenotype association mapping, Genetic interaction, Linear mixed model, 3D genomic interactions, Autism spectrum disorder

## Abstract

Biological interactions are prevalent in the functioning organisms. Correspondingly, statistical geneticists developed various models to identify genetic interactions through genotype-phenotype association mapping. The current standard protocols in practice test single variants or single regions (that contain multiple local variants) sequentially along the genome, followed by functional annotations that involve various aspects including interactions. The testing of genetic interactions upfront is rare in practice due to the burden of testing a huge number of combinations, which lead to the multiple-test problem and the risk of overfitting. In this work, we developed interaction-integrated linear mixed model (ILMM), a novel model that integrates a *priori* knowledge into linear mixed models. ILMM enables statistical integration of genetic interactions upfront and overcomes the problems associated with combination searching.

Three dimensional (3D) genomic interactions assessed by Hi-C experiments have led to unprecedented biological discoveries. However, the contribution of 3D genomic interactions to the genetic basis of complex diseases has yet to be quantified. Using 3D interacting regions as *a priori* information, we conducted both simulations and real data analysis to test ILMM. By applying ILMM to whole genome sequencing data for Autism Spectrum Disorders, or ASD (MSSNG) and transcriptome sequencing data (GTEx), we revealed the 3D-genetic basis of ASD and 3D-eQTLs for a substantial proportion of gene expression in brain tissues. Moreover, we have revealed a potential mechanism involving distal regulation between FOXP2 and DNMT3A conferring the risk of ASD.

Software is freely available in our GitHub: https://github.com/theLongLab/Jawamix5

## Introduction

Genotype-phenotype association mapping has revealed thousands of loci associated with complex traits. As genes function by various forms of interactions and no gene operates in a vacuum (Costanzo et al. 2016; Phillips 2008), it is possible that the discovered single-gene associations may represent only the tip of an iceberg of the genetic-basis of complex disorders. In Statistical Genetics, genetic interaction is defined as the non-linear effects between multiple loci (Baryshnikova et al. 2013). Although researchers have developed many statistical models aiming to discover the role of genetic interactions underlying complex disorders (Fang et al. 2019; Greene et al. 2015; Jansen et al. 2019a; Miguel-Escalada et al. 2019; Watson et al. 2019; Wen et al. 2016), single locus analyses such as single variant models (Kang et al. 2010) or approaches jointly analyze multiple local variants in a single regions (Wen et al. 2016; Wu et al. 2010) are still dominant in the practice of association mapping (Jansen et al. 2019b; Watson et al. 2019). This may be partly due to the difficulties of interpreting the large number of outcomes from interaction analyses as well as the problem of multiple-test and overfitting (Cordell 2009). In contrast, researchers frequently utilize information regarding interactions, such as chromatin status (Tak and Farnham 2015), transcriptional regulations (Gallagher and Chen-Plotkin 2018), and protein bindings (Mao et al. 2016) and others (Schork et al. 2013), from various databases (Gallagher and Chen-Plotkin 2018; Ward and Kellis 2012) as sources for downstream annotations of peaks from single-region association mappings. On the other front, many methods quantitatively integrate single-locus functional information in association studies were also developed (Kichaev et al. 2019; Lu et al. 2016; Pickrell 2014; Sveinbjornsson et al. 2016; Yang et al. 2017). Moreover, methods integrating known biological network information in association studies were also recently proposed (Carlin et al. 2019), although genetic principle (e.g., heritability) was omitted in building such models. To our best knowledge, there is no tool of association mapping that integrates *a priori*, however yet incomplete, knowledge of interactions of multiple genomic regions into the statistical genetic models.

In this work, we developed ILMM, Interaction-Integrated Linear Mixed Model, a novel tool integrating *a priori* knowledge of genetic interactions with a statistical test to map associations between interacting genetic regions and the phenotypic variations. By leveraging a linear mixed-model, ILMM only requires the knowledge of the potential genetic regions; ILMM can test the existence of joint effects when the actual mechanism of interaction is unknown (**Methods & Materials**). By integrating biological *a priori* knowledge into the statistical models, ILMM has two major advantages over state-of-the-art models. First, as mentioned above, models searching for potential combinations of genetic loci *de novo* may lead to an astronomical number of candidates (Hoh and Ott 2003). However, ILMM tests a controllable number of combinations based on prespecified interacting regions, therefore, significantly relieving the risk of overfitting and the burden of multiple-test corrections. Second, instead of conducting statistical tests and the interpretation of interactions sequentially, ILMM allows simultaneous modeling and quantitative assessment of *a priori* partial knowledge using genetic and phenotypic variations in the disease cohort. ILMM moves the efforts of collecting biological knowledge of interactions from downstream annotations to the upstream statistical tests. This is a meaningful innovation because the test P-values specifically quantify the strength of associations in terms of interactions as well as genetics rather than simply labelling marginal significance of a single gene as “found” or “not found” in the interactions databases.

The three-dimension (3D) chromatin structure is an important mechanism altering gene transcriptions that has been extensively studied for several years (Eres et al. 2019; Mah and Won 2019; Melo et al. 2020; Miguel-Escalada et al. 2019; Rao et al. 2014; Won et al. 2016). In addition to many biological insights revealed by the 3D structure of genomes, a fundamentally new view for statistical geneticists is that genes that are far apart when placed in the one-dimensional view could actually be spatially close to each other in 3D space when they function. This insight may bring a paradigm shift to statistical genetics, although more rigorous statistical analyses are required. A recent study suggested that the topologically associating domain (TAD) generally does not overlap with linkage disequilibrium (LD) block in large scale (Whalen and Pollard 2019). However, it is still unclear whether genetic interactions between the regions that are spatially close in a 3D domain exist and whether such interactions are functionally relevant to complex traits. Researchers have recently used 3D information in annotating peaks in association mappings (Fu et al. 2018; Giusti-Rodriguez and Sullivan 2019; Yu et al. 2019), however not integrated with association mapping models.

To illustrate the use of ILMM and further our understanding of the contribution of 3D genome structure to complex traits, we applied ILMM to multiple datasets of autism spectrum disorder (ASD) (Neale et al. 2012; Yuen et al. 2017) and gene expression data of 9 brain tissues and whole blood in the GTEx dataset (Aguet et al. 2017; Ardlie et al. 2015). In the analysis, we considered the interacting regions in 3D structure in brain tissues, assessed by Hi-C experiments (Rajarajan et al. 2018), as *a priori* information, and used them in the association mapping. We call this process 3D-Genome-wide association study, or 3D-GWAS. Indeed, we discovered interesting associations between 3D structure and ASD and expressions, which we called 3D-genetic basis of complex traits and 3D eQTLs, respectively. Additionally, we have identified substantial overlap between ASD and expressions in terms of 3D genetics, indicating pleiotropy effects shared by ASD and gene expressions. Through in-depth analysis of transcription factor binding motifs and protein docking, we have also revealed a mechanism underlying ASD that involves distal regulation between FOXP2 and DNMT3A.

This paper will explain the design intuition of ILMM, followed by its mathematical formulations. The simulations under various interaction mechanisms will be presented to demonstrate the universal power of ILMM which is robust to unknown mechanisms, contrasting to state-of-the-art alternatives. After that, analyses of real data and outcomes will be presented. Finally, limitations and future extensions will be discussed.

## Materials and Methods

### Design principle of ILMM

Multiple genetic regions can jointly alter the phenotype through various mechanisms, including epistasis (Bateson 1903; Mackay 2014), compensatory (Brown et al. 2010), heterogeneity (Madsen et al. 2011), or sometimes just additive (Madsen et al. 2011). Designing a general model to test interactions without knowing a specific mechanism could be tricky. A test that exhaustively verifies all potential known mechanisms would lead to substantial risk of overfitting and multiple-test burden; not to mention the lack of a complete list on all potential mechanisms. We took advantage of a specific angle underlying the linear mixed models; linear mixed models (LMM), although being called “linear”, can capture interactions implicitly. Although the pattern of genetic interactions could be complicated and largely unknown, the probability for two individuals carrying the same combinations of alleles at multiple genetic loci is proportional to the overall genetic similarity in these loci (e.g. identity by descendant). This similarity can be naturally captured by the genomic relationship matrix (GRM), which serves as the variance-covariance matrix in a multivariate normal distribution (MVN) of a random term in LMM.

Based on the above rationale, we designed ILMM by embedding the focal genetic regions into an LMM (**Fig. 1**). In this LMM, we have two random terms: one term employs the whole-genome GRM as the variance-covariance matrix in its MVN, while the other term employs a “interacting regional” GRM as its variance-covariance matrix. The interacting GRM is calculated using genetic variants in the regions that are suspected to have interactions, e.g., the two regions that are interacting in 3D space (revealed by a Hi-C experiment) (**Fig. 1**). Using such an LMM model, we aggregate the genetic variants in regions that are potentially interacting into the “regional” random term; and the rest contributions (i.e., other genes or population structure) are captured by the “global” random term using the whole-genome GRM.

**Figure 1.**
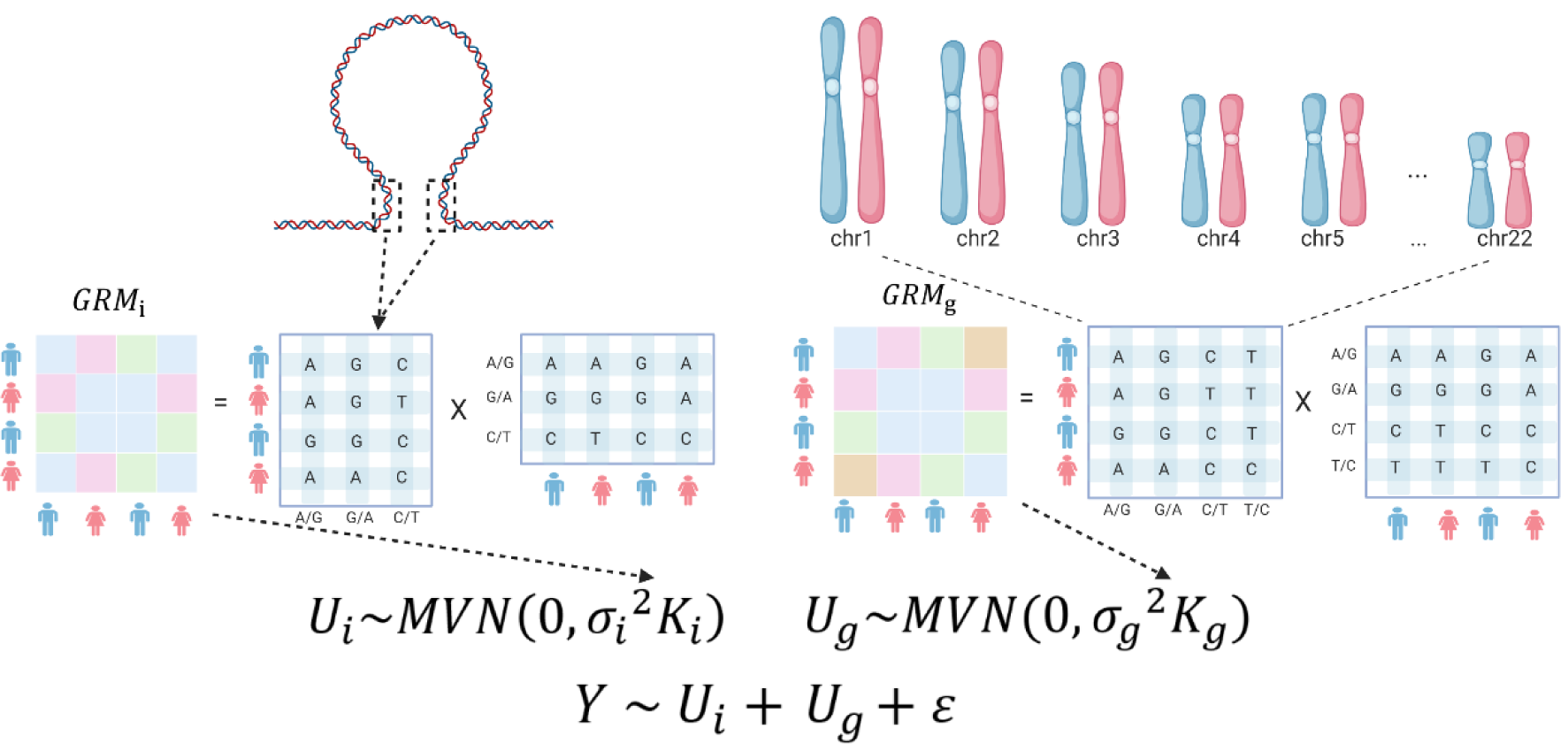
The protocol of ILMM. *GRM*_*i*_ refers to the genomic relationship matrix (GRM) calculated by the genotype present in the interacting regions. *GRM*_*g*_ refers to the GRM calculated by all genotype from the whole genome.

### Mathematical formulations

We use *Y* to denote the phenotype, *U*_*i*_ to denote the interaction term capturing the interacting regions, and *U*_*g*_ to denote the random term capturing the rest contribution. Our model then becomes:

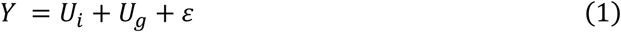

Where 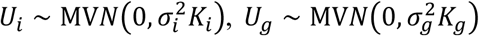, and 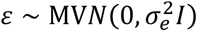.

where *I* isthe identity matrix, *K*_*g*_ is the GRM calculated using the whole genome variants: 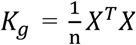, where *X* is the centralized and standardized genotype matrix and n is the number of variants in whole genome. Denoting the corresponding centralized and standardized genotype matrix in the *m*-th focal region (*m* = 1,2, …, *M*) as *X*_*m*_, and the combined genotype matrix *X*_*int*_ = (*X*_1, …,_ *X*_*m*_) then <u>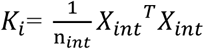</u>., where n_*int*_ is the total number of variants in all the *m* regions. The model is solved using an integration of de-correlation of the global GRM and the low-rank trick proposed by FaST-LMM (Lippert et al. 2011). The details of the mathematical derivations are in **Supplementary Materials**.

### Software implementation

ILMM is implemented as a function in our existing software Jawamix5 (Long et al. 2013; Xiong et al. 2019) that employs memory virtualization techniques (or “out-of-core” in computer science) based on HDF5 libraries. The software is scalable to very large genomic dataset that cannot be loaded into memory. The data file is stored in the disk with highly effective indexing so that the calculation is as fast as though the data were resided in the main memory (RAM).

### Procedure of simulations and type-I error adjustment

We thoroughly tested ILMM via simulations, contrasting to state-of-the-art alternatives. The control dataset of Wellcome Trust Case-Control Consortium (WTCCC) (Burton et al. 2007) were used as the template genotype (N=2,938).

To prepare *a priori* knowledge, we first collected a list of potential 3D-interacting pairs of regions, which has been reported by other researchers by conducting Hi-C experiments in brain tissues (Rajarajan et al. 2018; Watson et al. 2019; Won et al. 2016). Here, we utilized a Hi-C assessment in the developing brain which has 27,982 brain-specific paired 3D-interacting regions, measured from neurons derived from human induced pluripotent stem cells (hiPSCs) (Rajarajan et al. 2018). This dataset is available in the Synapse database (https://www.synapse.org/) with Synapse ID: syn12979149.

During each round of simulation, a pair of interacting regions was randomly selected. We then randomly selected 5 genetic variants in each region as causal to form the genetic contribution from the region. These 5 variants were modeled using a combination of “additive model” and “heterogeneity model”: the aggregated contribution from a region will be 0, 1, or 2 if there are 0, 1 or 2 alternate alleles in the 5 variants, reflecting an additive pattern. However, if there are more than 3 alternate alleles, the total contribution is still 2, reflecting heterogeneity pattern in which different mutations can cause phenotypic changes independently. Based on the regional contributions, the total genetic contribution to phenotypes were simulated using four mechanisms of joint contributions from multiple genetic regions; the models used are the additive model, epistasis model, heterogeneity model and compensatory model, defined in **Table 1**. Finally, the phenotype was simulated by adding genetic contribution with a random noise. We rescaled the genetic contribution to ensure that the phenotype indeed has the prespecified heritability.

**Table 1.** Definition of four different models used to simulate genetic component to phenotype (before adding random noise).

| **Additive** |  | Region A contribution |  |  |
| --- | --- | --- | --- | --- |
|  |  | 0 | 1 | 2 |
| Region B contribution | 0 | 0 | 1 | 2 |
|  | 1 | 1 | 2 | 3 |
|  | 2 | 2 | 3 | 4 |
| **Epistasis** |  | Region A contribution |  |  |
|  |  | 0 | 1 | 2 |
| Region B contribution | 0 | 0 | 0 | 0 |
|  | 1 | 0 | 1 | 1 |
|  | 2 | 0 | 1 | 1 |
| **Heterogeneity** |  | Region A contribution |  |  |
|  |  | 0 | 1 | 2 |
| Region B contribution | 0 | 0 | 1 | 1 |
|  | 1 | 1 | 1 | 1 |
|  | 2 | 1 | 1 | 1 |
| **Compensatory** |  | Region A contribution |  |  |
|  |  | 0 | 1 | 2 |
| Region B contribution | 0 | 0 | 1 | 2 |
|  | 1 | 1 | 0 | 1 |
|  | 2 | 2 | 1 | 0 |

ILMM was compared to EMMAX (Kang et al. 2010), the most frequently used tools for single-marker analysis, and SKAT (Wu et al. 2010; Wu et al. 2011), a popular method testing aggregated effects of genetic variants in a single region. We also tested ILMM against our own implementation of single-region-based mixed model, which is a specific case of ILMM when M=1 (i.e., the number of interacting regions is just one). We call this method “LOCAL”. The motivation of comparing ILMM to LOCAL is to quantify how much power gain ILMM will achieve when comparing with the single-region-based method with the same implementation.

To conduct a fair comparison, we simulated random phenotype (i.e., no genetic component) to adjust the Type-I-Errors (TIEs) to be the same. In particular, we ranked all the P-values calculated under the null distribution (i.e., using the random phenotype) and took the top 5% cut-off. If this cut-off is surrounding 0.05, we can conclude that the corresponding method’s TIE is under control. Also, this cut-off, after multiple-test correction, will be the cut-off to claim significance when calculating powers under alternative hypothesis. How the other tools were executed are detailed in **Supplementary Materials**.

### Genotype and phenotype data for 3D-GWAS

The *a priori* information of genetic regions suspected undergoing interactions, as stated in *Simulations*, is the 3D interaction data generated by other researchers using Hi-C experiments in brain tissues (Rajarajan et al. 2018).

The genotype and phenotype data are acquired from dbGaP and other genomic consortia. There are two ASD datasets: the first dataset is the influential MSSNG data (Yuen et al. 2017), containing 7,065 whole genome sequencing data; the second dataset contains 9,428 subjects (Neale et al. 2012) assessed by whole exome sequencing (phs000298.v4.p3.c1 & phs000298.v4.p3.c2). A more detailed description of the datasets is in **Supplementary Table S1**. We have conducted general quality control by removing variants with low MAF and derivation from HWE for association mapping but use the full dataset for downstream annotations.

Additionally, we applied ILMM to the gene expressions in brain tissues generated by the Genotype-Tissue Expression (GTEx) project (Aguet et al. 2017; Ardlie et al. 2015). The list of tissues and sample sizes are listed in **Supplementary Table S2**.

### Annotating 3D-GWAS outcomes by distal regulation

In this work, relating to a pair of interacting regions assessed by Hi-C experiments, distal regulation refers to the situation where a transcription factor (TF) binding to one region of the pair interacts with a gene located in the other region of the pair. The input of this analysis is a list of identified candidate genes (that are located in regions significantly associated with ASD assessed by ILMM P-values, e.g. DNMT3A). The output is a list of TFs (together with their binding motifs) regulating the candidate genes. To detect such regulations, we took two steps. First, we used the TF2DNA (Pujato et al. 2014) database to identify TFs which have been reported to interact with at least one of the candidate genes. These TFs, however, may or may not bind to the iterating regions. So, in the second step, we used the JASPAR (Khan et al. 2018) database to search for the binding sites (i.e., motifs) to filter out the TFs that do not bind to our candidate interacting regions. The TFs that have their binding sites located to the pairing regions will then be the output.

Next, we looked for genetic SNPs located at the binding sites (motifs) of these candidate TFs and further predicted the interactions between DNA and proteins (motif-TF complexes). This was achieved by utilizing HDock, a tool specifically designed to quantify the binding affinities between TF and motifs (Yan et al. 2020).

### Calculating LD between interacting regions associated with phenotype and expressions

The paired regions that are in genetic interactions may be susceptible to have high linkage disequilibrium (LD), even though they may be far away from each other. For selected pairs of regions, we computed their LD in terms of both D’ and r^2^ using genotype data from the 1000 Genomes Project (Altshuler et al. 2015). Since the standard LD is defined between two genetic variants, we calculated the pairwise LD between all variants in one region and all variants in the other region and considered their average as the LD between two regions. To contrast the LDs between interacting regions with the background, i.e., non-interacting regions, we formed a null distribution by calculating LDs between randomly selected pairs of regions with the same sizes and between-region distance for 1000 times. These 1000 average LD values formed a null distribution. Then the standings of actual LDs of interacting regions will be ranked in the null distribution as an assessment of whether they are significantly high.

## Results

### Simulations

The type-I-errors of all the four competing methods were generally under control, although ILMM and LOCAL are slightly different to the supposed value of 0.05 (**Supplementary Table S3**). In particular, ILMM was more conservative (with a top 5% cut-off being 0.0713). Using the adjusted critical values ensuring type-I-errors to be 0.05, the fairness of power comparison was guaranteed. QQ-plot for the ILMM P-values under null hypothesis showed that the expected P-values are generally equal to observed P-values, although they were slightly under the diagonal (**Supplementary Fig. S1**), consistent to the slightly conservative type-I error.

As described in **Methods and Materials**, we compared the power of ILMM with other three methods: SKAT, EMMAX and LOCAL. The power is defined as the number of rounds that the corresponding method significantly identified the simulated pairs of regions. In the present setting of pairs of regions, there are two regions to be detected. For ILMM, naturally it will detect both regions in a single test; however, for the other tests, the criteria of defining “success” could be detecting at least one region or detecting both regions. Here we used “(1)” to indicate the criteria of detecting at least one region, and “(2)” for the criteria of detecting both interacting regions. SKAT and LOCAL are naturally region-based, and we set up the window size of 5kb, which is the average of the length of one side of the pairs of interacting regions. For EMMAX, which is based on a single marker test, we claimed success of a region as long as there is at least one genetic variant significantly associated with the phenotype. The statistical significance of a particular test for a focal method is defined by its P-value smaller than the corresponding cut-off observed in the previous simulations to adjust type-I-error (**Supplementary Table S3)** divided by the number of tests (i.e., Bonferroni correction (Noble 2009)). Note that different methods had different numbers of tests. For SKAT and LOCAL, the number of tests was 1200K (=3 × 10^9^/2500), which was the total number of tiling windows across the genome. For ILMM, it was 27,982, the total number of Hi-C assessed spatial-interacting regions. For EMMAX, it was the number of SNPs of the WTCCC array, which was 386,469.

The outcome is depicted in **Fig. 2**. Evidently, the other methods had very small powers when the criterion is to detect both interacting regions. This is consistent with the motivation that testing multiple interacting regions together will substantially improve power. Additionally, ILMM outperforms all the methods with the criteria of identifying at least one region. This implied that the current standard protocol, which discovers an associated region in single-region tests followed by database-search-based annotations, is suboptimal compared to the interaction-integrated test.

**Figure 2.**
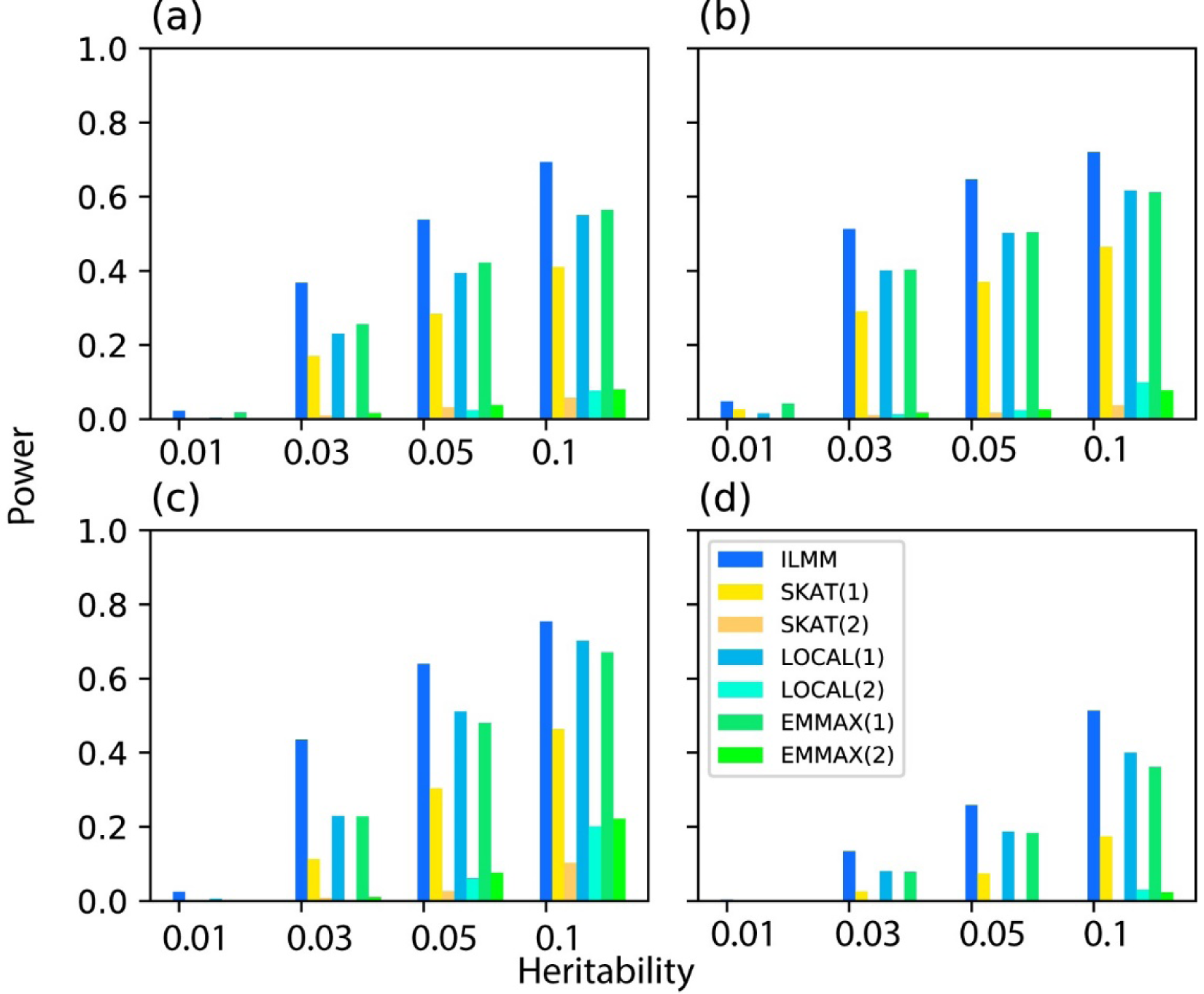
Statistical power (y-axis) of four methods under four interacting models with various regional trait heritability (x-axis). (a): Additive model, (b): Epistasis model, (c): Heterogeneity model, (d): Compensatory model

### 3D-genetic basis of Autism Spectrum Disorder

Using the potential 3D interacting regions revealed by Hi-C experiments in neurons induced by human iPSC brains (Rajarajan et al. 2018) as *a priori* knowledge, we applied ILMM to the MSSNG whole-genome sequencing data (Yuen et al. 2017) to identify 3D-genetic basis of ASD. As important follow-ups, we carried out annotations for the statistically significant genes from various aspects including enrichment analysis for pathway and gene ontology, as well as functional annotation towards distal regulations.

### Statistically significant genes

By applying ILMM to MSSNG dataset, we identified 1,164 pairs of regions that are significantly associated with ASD (whole genome FDR =0.05). There are 1,445 genes located in these significant regions (**Supplementary Table S4**). As a comparison, we also applied SKAT (Wu et al. 2010; Wu et al. 2011) to this dataset with sliding windows of 5kb and revealed 32,682 significant regions (with the same criteria of whole-genome FDR =0.05), containing 6,322 genes, among which 663 genes (10.4%) also have been identified by ILMM method (**Supplementary Table S5**). This comparison indicates that majority genes still impact ASD by marginal effects, in which around 10.4% may be involved in 3D-interactions. On the other hand, by directly integrating interactions in a statistical model, ILMM identified around 2.1 (= 1,445 / 663) times genes than a single region-based model.

To carry out functional enrichment analyses for the 1,445 potential genes located in significant regions, we utilized an R package named clusterProfiler (Yu et al. 2012) which quantifies the level of enrichment of a gene set with regards to various functional annotations. Using clusterProfiler, we performed both pathway enrichment analysis based on Kyoto Encyclopedia of Genes and Genomes (KEGG) (Kanehisa et al. 2002) and gene ontology (GO) (Ashburner et al. 2000) enrichment. The top 20 KEGG significantly enriched pathways were reported (**Fig. 3a**). Among them, Huntington disease, which belongs to the nervous system category, was the pathway that covered the highest proportion of genes. Notch signaling pathway, which is central to a wide range of development processes in human organs (Lasky and Wu 2005) was also among the most significantly enriched pathways. Notably, the Glutamatergic synapse pathway, which is labelled by KEGG to be associated with ASD, was also present in our results. Several additional pathways, such as antigen processing and presentation, SNARE interactions in vesicular transport and base excision repair, were also reported to be related with ASD (Bennabi et al. 2018; Castermans et al. 2010; Markkanen et al. 2016).

**Figure 3.**
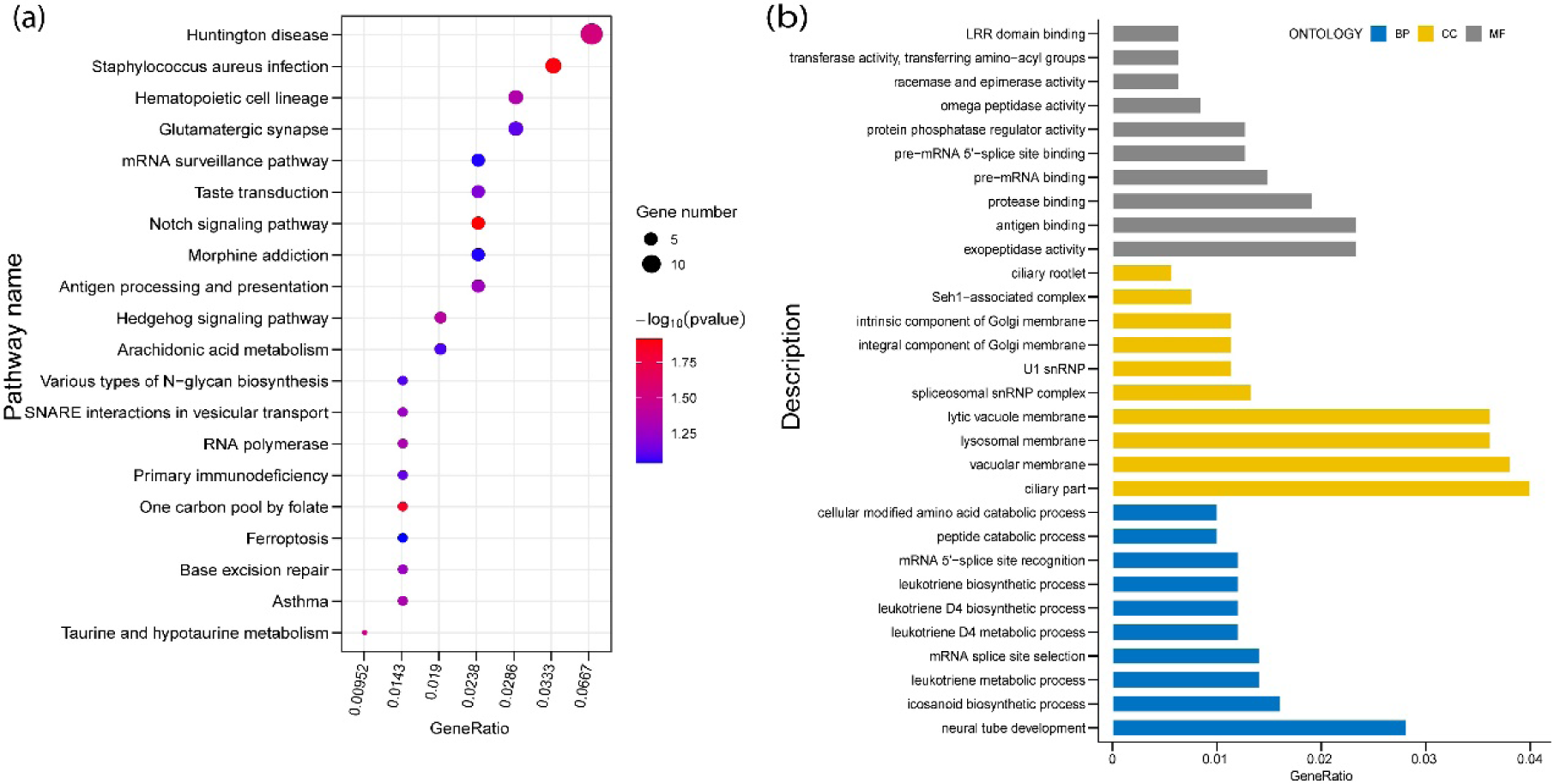
KEGG and GO enrichment analysis of significant genes. (a): Top 20 KEGG pathways (ranked by p-value). Gene ratio (x-axis) is the percentage of significant genes over the total genes in a given pathway. (b): Top 10 (ranked by p-value) GO terms of three categories (BP: biology process, CC: cell component, MF: molecular function). Gene ratio (x-axis) is the percentage of the number of genes present in this GO term over the total number of genes in this category.

The GO analysis was applied to three ontologies: biological process (BP), cellular component (CC), and molecular function (MF); and the top 10 significant GO terms are depicted (**Fig. 3b**). Among the top 10 highly enriched BP terms, four of them are leukotriene associated, i.e., leukotriene metabolic process, leukotriene D4 metabolic process, leukotriene D4 biosynthetic process, and leukotriene biosynthetic process. The elevated levels of leukotrienes have been reported in autistic patients in several studies (El-Ansary and Al-Ayadhi 2012; Theoharides et al. 2016). In addition, leukotriene can be used as a biomarker for the early diagnostic of autistic patients (El-Ansary and Al-Ayadhi 2012; Qasem et al. 2016). Among the significant CC terms, U1 snRNP (small nuclear ribonucleoprotein) is the most significant cellular component (P-value = 1.06 × 10^−4^) and is one of the 9 snRNA blood signatures boosting diagnostic accuracy for ASD in clinical practice(Zhou et al. 2019). For MF, the most significant enrichment was pre-mRNA 5’-splice site binding (P-value = 2.23 × 10^-5^), consistent with the report of the important role of alternative splicing of mRNA in ASD blood (Stamova et al. 2013). The above analyses for KEGG and GO enrichment showed that, at the gene-set level, our discoveries generally align to reported ASD pathology.

In addition to the enrichment analyses, we conducted further annotations of significant results to figure out a hypothetical novel mechanism underlying ASD. To narrow down candidates, we first searched the SFARI database, an established repository for existing ASD genes (Abrahams et al. 2013), and found that 49 ILMM-identified genes in are in SFARI. The proportion that is in SFARI, 49/1,445 = 3.4%, is relatively low. This might be due to that firmly verified ASD genes are limited. Indeed, for the established method for single region analysis, SKAT, the corresponding ratio is also 217/6,322 = 3.4%. These 49 genes include 12 genes scored as 1, which are known as high-confident ASD genes in SFARI (**Table 2**).

**Table 2.**
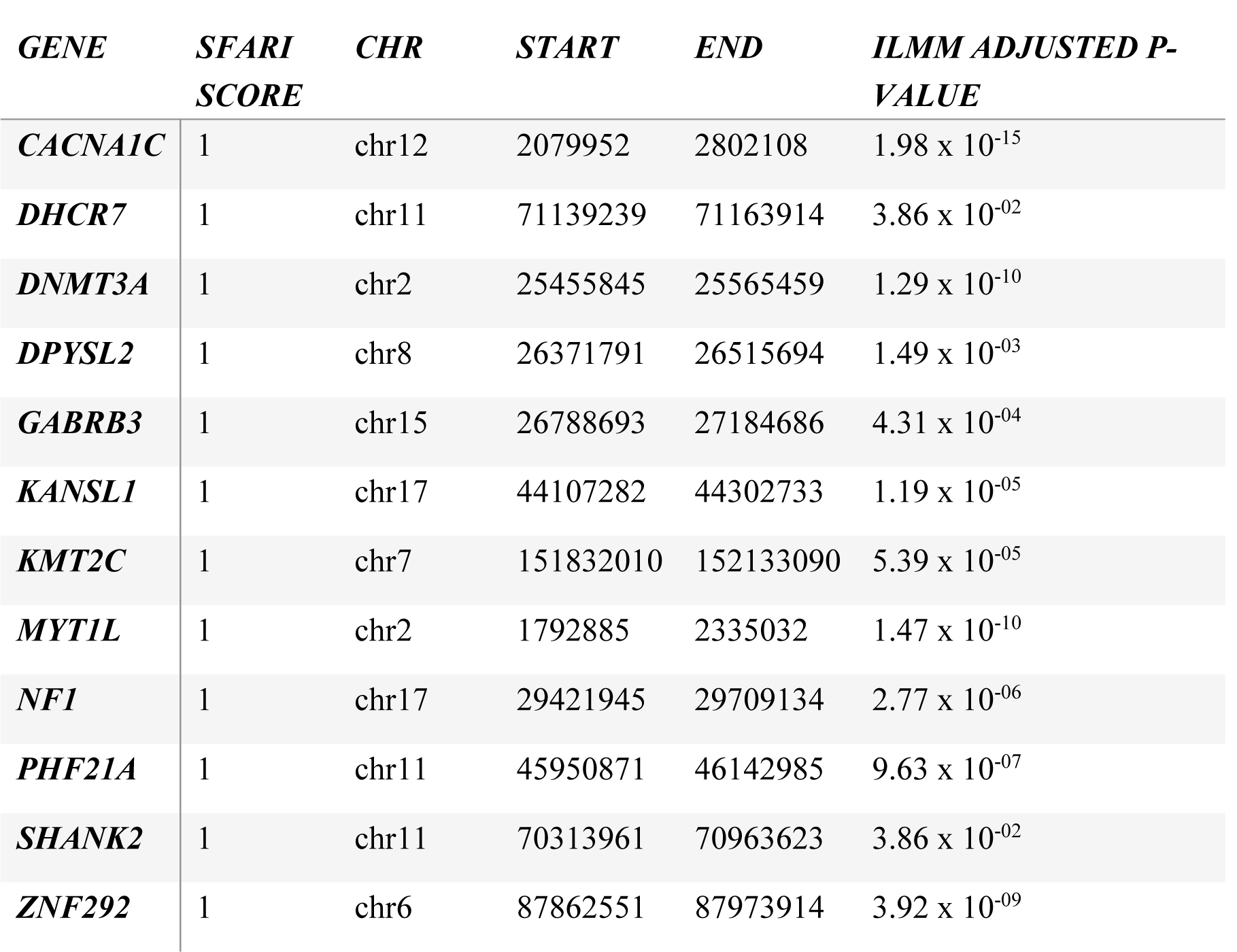
Significant genes identified by ILMM which are also reported as high confident ASD genes in the SFARI database. An FDR of 0.05 is used to adjust P-values.

Using the TF2DNA + JASPAR pipeline (described in **Materials and Methods**), we generated the distal regulatory TFs for the above score-1 genes. As the MSSNG dataset is whole-genome sequencing that provides all polymorphisms, we were enabled to characterize the consequence of a genetic mutation located in a motif to the binding complex by using HDock (Yan et al. 2020). In particular, reference and mutant binding motifs as well as the TFs (represented by their protein data bank ID, or PDB) were submitted to the HDock server. We found that (1) FOXP2 is a TF to the gene DNMT3A (a score-1 candidate gene in **Table 2**), and (2) a single nucleotide polymorphism, or SNP (NC_000002.11: g.133032477T>C) on the binding motif of FOXP2 (ATTGTT**T**TATT), will affect FOXP2 binding affinity. More specifically, on the structure of FOXP2, there is a positively charged interface (J:542R, J:543R, K:553R, K:554H, K:549K, K:583R, where J and K are protein chain index) in the minimum energy protein-ligand complex around the reference allele T (**Fig. 4a**). According to a previous study (Luscombe et al. 2001), because thymine has the highest acidities among the four nucleotides, thymine (reference allele) preferentially interacts with arginine (R), histidine (H), and lysine (K) than cytosine (mutant allele). Thus, thymine can reduce the binding energy by forming stable protein-DNA electric fields (Luscombe et al. 2001). Consistent to this interpretation, the docking score for wild type motif is -362.00, in contrast to the docking score for the mutated motif (ATTGTT**C**TATT) being -330.00, which has higher binding energy with FOXP2. Together, this suggested that the mutation (g.133032477T>C) reduces the binding affinity of FOXP2 to its motif. As a result, DNMT3A, which is regulated distally by FOXP2 through chromatin loop, may have diminished expression (**Fig. 4b**).

**Figure 4.**
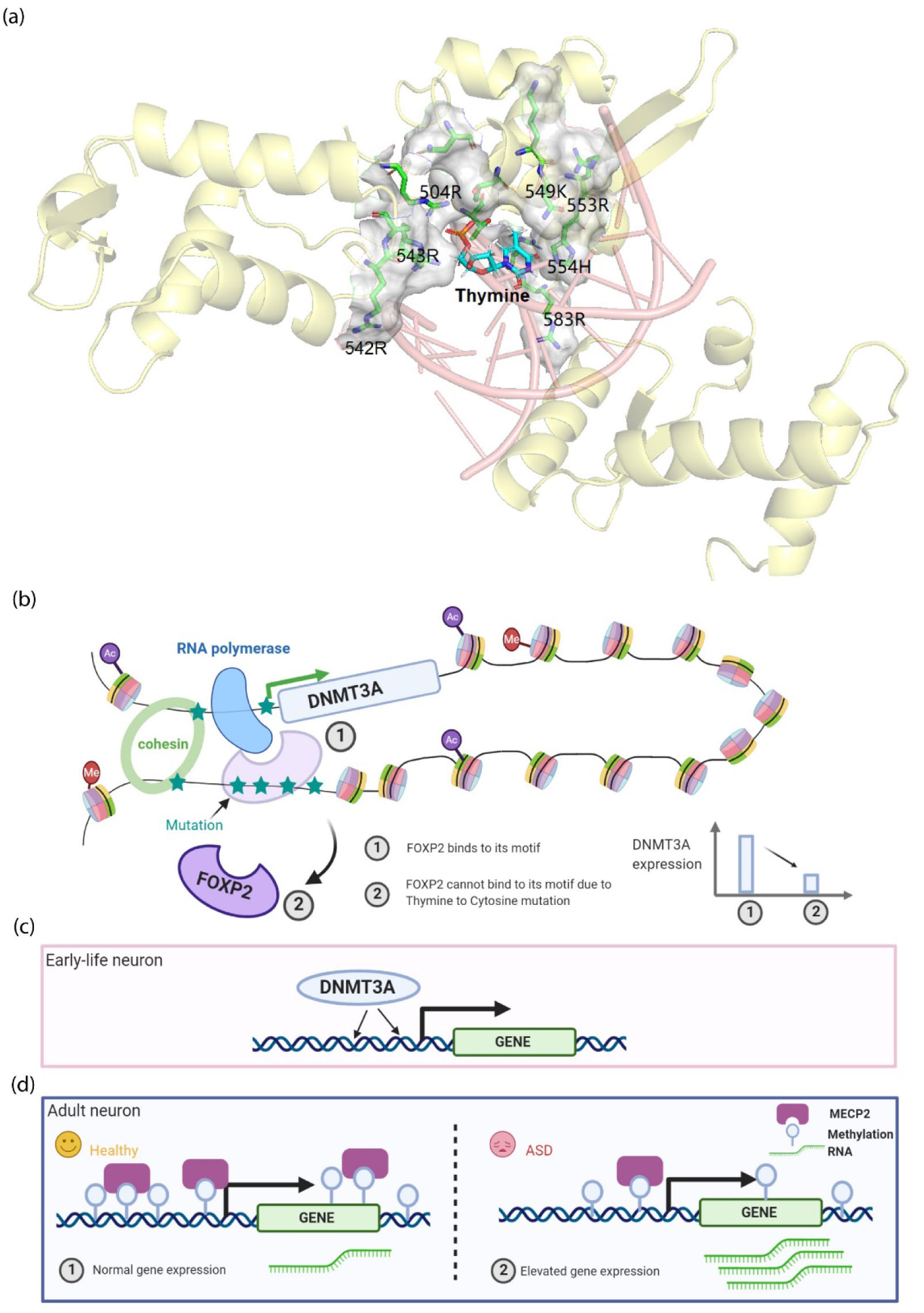
Interaction between FOXP2 and its motif for distal regulation on DNMT3A. (a) Thymine (reference allele) on the motif and its nearby positively charged interface of FOXP2 visualized by PyMOL (DeLano 2002). (b): The single nucleotide mutation on FOXP2 motif is likely to reduce the binding affinity of FOXP2 which may decrease the gene expression of DNMT3A through distal regulation. (c): A gene present in early-life neuron and DNMT3A deposits methylation on its CA sequences. (d): The low level of DNMT3A across the neuron development process may cause hypomethylation of many genes resulting in elevated expression of these genes in adult neurons.

DNMT3A encodes an enzyme named DNA methyltransferase 3 alpha, which is involved in DNA methylation and plays a crucial role in epigenetic regulation in cells. In particular, DNMT3A binds preferentially to intergenic regions and across transcribed regions of genes, which primarily induces methylated CA sequences (mCA) in the early-life neuron (Stroud et al. 2017) (**Fig. 4c**). These mCA functions act as landmarks for MECP2, whose function is to restrain gene expression in the maturing brain (**Fig. 4d**). As such, DNMT3A, mCA, and MECP2 coordinate to precisely tune gene expressions that are crucial for normal brain development and function (Stroud et al. 2017). The hypomethylation in adult neurons caused by insufficient DNMT3A at their early life will lead to overexpression of many genes that lead to risk of ASD (Stroud et al. 2017) (**Fig. 4d**). Indeed, disruption of the cooperation either through DNMT3A or MECP2 has been reported to cause Rett syndrome, a severe neurological disorder with features of autism. (Chahrour and Zoghbi 2007; Gabel et al. 2015). Additionally, mutations on DNMT3A have been widely reported in ASD (Alex et al. 2019; Yokoi et al. 2020) as well as those with intellectual disabilities (Tatton-Brown et al. 2014). As such, our proposed mechanism is that mutations in the region on or surrounding DNMT3A in conjunction with mutations of FOXP2 binding sites jointly confer the risk of ASD.

In summary, our in-depth analysis including ILMM-based genetic mapping, functional annotation, motif search, and protein docking has jointly revealed a plausible mechanism for ASD: the SNV (g.133032477T>C), presenting in one of FOXP2 motifs, may lead to decreased gene expression of DNMT3A through distal regulation. The low level of DNMT3A may further cause the hypomethylation of CA, which reduces the recruitment of MECP2 and results in the increased expression of some genes, causing higher risk to ASD. This regulation through 3D interactions may jointly confer the risk of ASD with local mutations surrounding DNMT3A.

### 3D-cis eQTL in brain tissues

EQTLs are the genetic mutations associated with gene expression. Analog to the above 3D-GWAS that discovers 3D-genetic basis of complex traits, here we aimed to extend the concept of eQTL to spatially interacting regions using the list of interacting regions assessed by Hi-C experiments (that were used above). To keep this first attempt simple, we only looked at the eQTLs in *cis*, which means the gene body is located in or surrounding one of the interacting regions (20,000 base pair upstream or downstream of that gene). Such 3D-cis eQTLs may be deemed as *trans* in standard analysis (with 1D genome) but are considered as *cis* here as we hypothesized that the regulatory mutations are spatially nearby in the 3D domain. We applied ILMM on 10 tissues in GTEx datasets including 9 brain tissues and the whole blood to identify interacting regions functioning as 3D-cis eQTLs. First, we selected genes with high variance (variance >= 10 in RPKM value), which led to around five thousand genes for each tissue. Then we performed eQTL mapping between expressions of these selected genes and genotypes using ILMM to uncover paired regions that are significantly associated with gene expressions. Based on a cut-off of P-value lower than 0.05 after FDR correction (Noble 2009), we discovered hundreds of significant 3D-cis eQTL from these 10 tissues. All results are listed in **Supplementary Table S6-15**. To assess the proportion of genes that are were able to detect 3D-cis eQTLs, we calculated the number of genes that are located in (or around) a pair of interacting regions and the number of genes for which we indeed identified 3D-cis eQTLs. It is observed that 3D-cis eQTL accounts for a small proportion (3% - 8%) of genes (**Fig. 5a**). This indicates that 3D-cis eQTLs do exist, however, they are not dominant for gene regulations.

**Figure 5.**
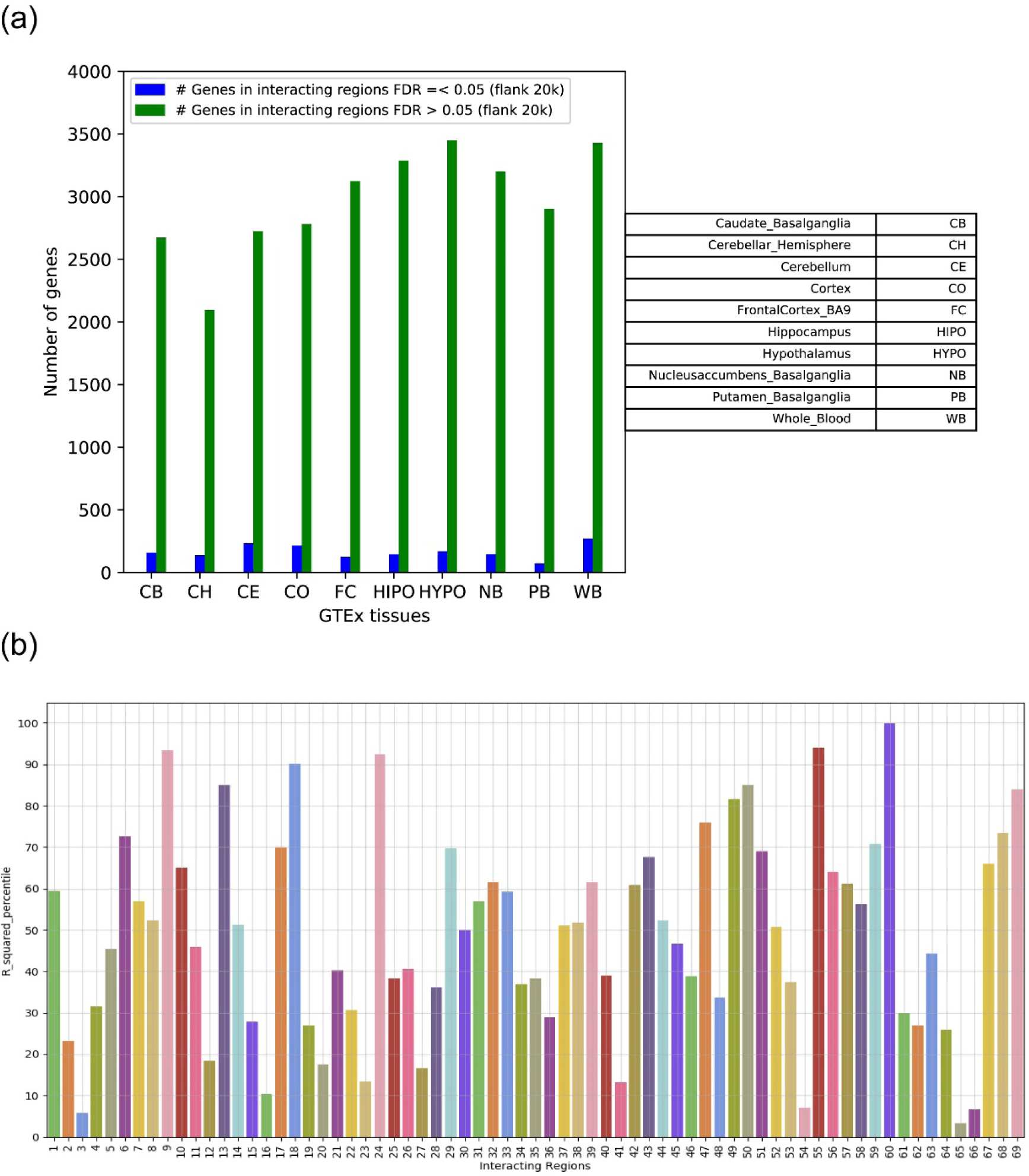
3D-cis eQTL analysis. (a) Number of genes for which we can and cannot identify 3D-cis eQTLs. (b) LD (r^2^) value for 69 interacting regions associated with both ASD and gene expressions. X-axis: coordinate of one interacting region (details in Supplementary Table S15). Y-axis: percentile of LD value among background (higher percentile indicates stronger LD in interacting regions).

We checked whether the 3D interacting regions are in higher linkage disequilibrium (LD) compared to background (**Materials and Methods**). To narrow down the candidates, we selected only the pairs of regions that are significantly associated with both ASD and gene expressions. In total, there were 69 paired regions as the 3D co-localized loci for both ASD and eQTL (**Supplementary Table S16**). We then calculated the average r^2^ and D’ between the two regions and contrasted them to the background (**Materials & Methods**). The outcome is depicted in **Fig. 5b & Supplementary Fig. S2**, showing that some of the regions indeed experienced significantly higher LD than the background. However, many regions did not. This showed that the 3D interactions may be very weak to the extent of not being able to impact the LD between regions, an observation consistent with the recent reported LD study at a larger scale (Whalen and Pollard 2019).

## Conclusion & Discussion

By applying ILMM to GTEx data, we identified 3D-cis eQTLs for only 3% - 8% genes in the brain tissues. This indicated that such 3D-genetic basis of expressions is not dominant, compared with standard cis eQTLs based on the 1D view of a genome. However, this may be due to our current limitation of accurately pinpointing the 3D interacting regions. Furthermore, the genetic interaction may not be limited to chromatin conformations, and other sources of interactions may also contribute as well. Therefore, there is potentially substantial room to utilize ILMM in practice.

As the GO and KEGG dataset may not be specifically designed for ASD, based on our hypothesis that ASD might be mediated by communication disorders, we specifically searched for annotations of genes association with hearing deficiencies using DisGeNET (Pinero et al. 2020). In total 41 genes that have been reported to be associated with Sensorineural Hearing Loss (disorder) or Nonsyndromic Deafness. Notably, 8 genes among these 41 genes were reported to be associated with ASD (**Supplementary Table S17**). Starting from these genes, further exploration to the functional mechanism of ASD medicated by communication disorders will be an interesting future work.

In addition to the whole genome data from MSSNG, we have also applied ILMM together with the same 3D interacting regions (Rajarajan et al. 2018) to a whole exome sequencing (WES) dataset containing 7,766 individuals (4944 control and 2822 ASD cases, dbGaP ID: phs000298.v4.p3.c1), which yielded no significant results. In contrast, applying a single-region tool with sliding windows of 5kb and 25kb led to the discovery of several significant regions (**Supplementary Table S18)**. This outcome indicates that the 3D genetic interactions are generally distributed in non-genic regions and may need sequencing data to analyze.

In summary, we developed a novel statistical method, ILMM, that integrates *a priori* knowledge of genetic interaction with linear mixed models to statistically identify genetic interactions. To demonstrate its use in practise, we used 3D chromatin conformation information assessed by Hi-C experiments as example *a priori* of potentially interacting regions. Using this list of paired regions, we applied ILMM to the whole-genome sequencing data generated by MSSNG and revealed the 3D genetic basis of ASD. Additionally, we also applied ILMM to transcriptome data generated by the GTEx consortium. The real data analysis revealed substantial insights into the 3D-genetic basis of both complex traits and expressions. Specifically, we formed a hypothetical mechanism including both local mutations and a motif impacting distal regulation that jointly confer the risk of ASD. This novel statistical method offers a complementary protocol to the standard practice that conducts association mappings for single regions followed by annotations and co-localization analyses. Additionally, compared with pure statistical methods searching for interactions *de novo*, ILMM does not suffer from the problem of choosing or optimizing the size of candidate genetic regions as the *a priori* knowledge will naturally offer such information. Therefore, ILMM further reduces the burden of multiple-test and risk of overfitting. Finally, the outcome of ILMM is naturally interpretable as the potential annotation, e.g., 3D-interactions, has been built in. We expect ILMM will be broadly used in practise to discover novel genetic interactions.

## Declaration

### Availability of data and material

All datasets used in this study are available in the following URL through application.

1. WTCCC data: https://www.wtccc.org.uk/info/access_to_data_samples.html
2. MSSNG data: https://www.mss.ng/
3. phs000298.v4.p3: https://www.ncbi.nlm.nih.gov/projects/gap/cgi-bin/study.cgi?study_id=phs000298.v4.p3
4. Genotype-Tissue Expression (GTEx) : https://www.ncbi.nlm.nih.gov/projects/gap/cgi-bin/study.cgi?study_id=phs000424.v8.p2
5. Brain Hi-C interacting regions: https://www.synapse.org/#!Synapse:syn12979149

### Code availability

Software is freely available in our GitHub: https://github.com/theLongLab/Jawamix5

### Authors’ contributions

[Q.L.1 = Qing Li and Q.L.2 = Quan Long]

Conceived the study: Q.L.2 and J.Y.; Developed the software: Q.L.2; Simulations: Q.L.1; Real data analysis: Q.L.1, C.C., and D.P.; Contributed to the analysis: J.H, X.C., F.A., and A.H.; Contributed to the interpretation of the data: B.A. and J.Y. Wrote the manuscript: Q.L.2, Q.L.1, and C.C. with the input of all authors.

