## Supplementary_notes for "Statistical model integrating interactions into genotype-phenotype association mapping: an application to reveal 3D-genetic basis underlying Autism"

### ***Mathematical formulations***

Our model to be solved is:

$$Y = u_i + u_g + \varepsilon \tag{1}$$

Where $u_i \sim N(0, \sigma_i^2 K_i)$, $u_g \sim N(0, \sigma_g^2 K_g)$, and $\varepsilon \sim N(0, \sigma_e^2 I)$.

**Step 1: estimating the genetic variance component $\sigma_g^2$**

First, we controlled the population structure through solving $\sigma_g^2$ to remove the correlation between individuals. We use $Y_c$, a centered $Y$, to regress on the random term $u_g$ and errors $\varepsilon$ as follows:

$$Y_c = u_g + \varepsilon \tag{2}$$

where $Y_c = Y - \bar{Y}$, $\bar{Y}$ is the average of $Y$, $u_g \sim N(0, \sigma_g^2 K_g)$, and $\varepsilon \sim N(0, \sigma_e^2 I)$. The eigen-decomposition of $K_g$ is $K_g = U_x S_x U_x^{-1}$, where $S_x = \begin{bmatrix} \lambda_{x1} & \cdots & 0 \\ \vdots & \ddots & \vdots \\ 0 & \cdots & \lambda_{xn} \end{bmatrix}$ is a matrix of eigenvalues. A new parameter $\delta = \frac{\sigma_e^2}{\sigma_g^2}$ is defined so that $Y_c \sim MVN\left(0, \sigma_g^2 (K_g + \delta I)\right)$. The estimated variance components for $\sigma_g^2$, $\sigma_e^2$ are therefore written in the following equations.

$$\hat{\sigma}_g^2 = \frac{1}{n} \sum_{i=1}^{n} \frac{V_{xi}^2}{(\hat{\delta} + \lambda_{xi})} \tag{3}$$

$$\hat{\sigma}_e^2 = \frac{\hat{\delta}}{n} \sum_{i=1}^{n} \frac{V_{xi}^2}{(\hat{\delta} + \lambda_{xi})} \tag{4}$$

Where $V_x = {U_x}^T Y_c$ , $\lambda_{xi}$ is the eigen value, $\hat{\delta}$ can be estimated by solving the non-linear equation below through Newton-Raphson method.

$$\sum_{i=1}^{n} \left[ \frac{n V_{xi}^2}{\left( \sum_{i=1}^{n} \frac{V_{xi}^2}{(\delta + \lambda_{xi})} \right) (\delta + \lambda_{xi})^2} - \frac{1}{(\delta + \lambda_{xi})} \right] = 0 \tag{5}$$

In Newton-Raphson method, we let $g(\delta_n) = \sum_{i=1}^{n}\left[\frac{nV_{ij}^2}{\left(\sum_{i=1}^{n}\frac{v_{xi}^2}{(\delta_n+\lambda_{xi})}\right)(\delta_n+\lambda_{xi})^2} - \frac{1}{(\delta_n+\lambda_{xi})}\right]$,

then by repeating the process: $\delta_{n+1} = \delta_n - \frac{g(\delta_n)}{g'(\delta_n)}$, until $|\delta_{n+1} - \delta_n| \leq 10^{-6}$, we can approximately solve the equation, yielding an estimate of $\delta$ (the variance ratio). Then we can calculate the estimates of $\hat{\sigma}_g^2$ and $\hat{\sigma}_e^2$. Then the decorrelation matrix $\widehat{D}_x$ can be formed:

$$\widehat{D}_x = \left(\hat{\sigma}_g^2 S_x + \hat{\sigma}_e^2 I\right)^{-\frac{1}{2}} U_x^T \quad (6)$$

**Proof of the soundness of the decorrelation procedure**

In equation (2) $Y_c = u_g + \varepsilon$

$$\begin{aligned} Var(Y_c) &= \sigma_g^2 K_g + \sigma_e^2 I \\ &= \sigma_g^2 U_x S_x U_x^T + \sigma_e^2 U_x U_x^T \\ &= U_x\left(\sigma_g^2 S_x + \sigma_e^2 I\right) U_x^T \end{aligned}$$

The following derivation justifies that $D_x = \left(\sigma_g^2 S_x + \sigma_e^2 I\right)^{-\frac{1}{2}} U_x^T$ will lead to the desired property that $Var(D_x Y) = I$.

$$\begin{aligned} Var(D_x Y) &= D_x Var(Y) D_x^T \\ &= D_x U\left(\sigma_g^2 S_x + \sigma_e^2 I\right) U_x^T D_x^T \\ &= \left(\sigma_g^2 S_x + \sigma_e^2 I\right)^{-\frac{1}{2}} U_x^T U_x \left(\sigma_g^2 S_x + \sigma_e^2 I\right) U_x^T \left(\left(\sigma_g^2 S_x + \sigma_e^2 I\right)^{-\frac{1}{2}} U_x^T\right)^T \\ &= \left(\sigma_g^2 S_x + \sigma_e^2 I\right)^{-\frac{1}{2}} U_x^T U_x \left(\sigma_g^2 S_x + \sigma_e^2 I\right) U_x^T U_x \left(\sigma_g^2 S_x + \sigma_e^2 I\right)^{-\frac{1}{2}} \\ &= \left(\sigma_g^2 S_x + \sigma_e^2 I\right)^{-\frac{1}{2}} I \left(\sigma_g^2 S_x + \sigma_e^2 I\right) I \left(\sigma_g^2 S_x + \sigma_e^2 I\right)^{-\frac{1}{2}} \\ &= \left(\sigma_g^2 S_x + \sigma_e^2 I\right)^0 \\ &= I \end{aligned}$$

Hence multiplying the decorrelation matric $D_x$ to $Y$ can control the population stratification by removing the correlation between individuals.

**Step 2: solving the local variance component $\sigma_l^2$**

After getting the decorrelation matrix $\widehat{D}_x$ from step 1, we applied this matrix to $Y_c$, and get $Y_c^* = \widehat{D}_x Y_c$. So, equation (1) can be reformat to equation (7) as below, where

$u_i \sim N(0, \sigma_i^2 K_i)$. The next step was to solve $\sigma_i^2$ using low-rank trick proposed by FaST-LMM.

$$Y_c^* = u_i + \varepsilon \quad (7)$$

For details, please refer to the original paper of FaST-LMM (Lippert et al. 2011).

### ***Details of running ILMM, LOCAL, EMMAX, SKAT***

ILMM, LOCAL, and EMMAX (Kang et al. 2010) methods are all implemented in Jawamix5 (Long et al. 2013; Xiong et al. 2019). More details can be found in the user manual in the GitHub (https://github.com/theLongLab/Jawamix5) for reference.

1. Convert genotype file from .csv format to .hdf5 format
    a. Command line: java – Xmx4g –jar /path/to/jawamix5.jar import -ig genotype.csv -o genotype.hdf5
    b. Parameters:
        i. -ig: input genotype file in plain text (.CSV format)
        ii. -o: output in HDF5 in format
    c. Input file: genotype.csv
    d. Output file: genotype.hdf5
2. Generate the genetic relationship matrices (GRM) based on input genotype file
    a. Command line: java –Xmx4g –jar /path/to/jawamix5.jar kinship -ig genotype.hdf5 -o genotype.kin
    b. Parameters:
        i. -ig: input genotype file in HDF5 format
        ii. -o: the output file prefix
    c. Input file: genotype.hdf5
    d. Output files:
        i. genotype.kin.rescaled.IBS
3. Run ILMM method
    a. Command line: java –Xmx4g –jar /path/to/jawamix5.jar compound -ig genotype.hdf5 -ip phenotype.tsv -o ./ILMM_res/ -ik_g genotype.kin.rescaled.IBS -ic hic_info.txt
    b. Parameters:
        i. -ig: input genotype file in HDF5 format
        ii. -ip: phenotype file
        iii. -o: output folder
        iv. -ik_g: the global genetic relationship matrices file

        v. -ic: input regions
    c. Input files:
        i. genotype.hdf5
        ii. phenotype.tsv
        iii. genotype.kin.rescaled.IBS
        iv. hic_info.txt (three columns separated by tab, an example listed below)
            #header: Index   Region1(chr; start; end)   Region2
            #content: C0   1;840000;850000   1;890000;900000
    d. Output file:
        i. ./ILMM_res/xxx.csv
4. Run LOCAL method
    a. Command line: java–Xmx4g –jar /path/to/jawamix5.jar local -ig genotype.hdf5 -ip phenotype.tsv -o ./LOCAL_res/ -ik_g genotype.kin.rescaled.IBS -w 5000
    b. Parameters:
        i. -ig: input genotype file in HDF5 format
        ii. -ip: phenotype file
        iii. -o: output folder
        iv. -w: tiling window size
        v. -ik_g: the global genetic relationship matrices file
    c. Input file:
        i. genotype.hdf5
        ii. phenotype.tsv
        iii. genotype.kin.rescaled.IBS
    d. Output file:
        i. ./LOCAL_res/xxx.csv
5. Run EMMAX method
    a. Command line: java–Xmx4g –jar /path/to/jawamix5.jar emmax -ig genotype.hdf5 -ip phenotype.tsv -o ./EMMAX_res/ -ik genotype.kin.rescaled.IBS -p 0.05
    b. Parameters:
        i. -ig: input genotype file in HDF5 format
        ii. -ip: phenotype file
        iii. -o: output folder
        iv. -ik: genetic relationship matrices file generated by function “kinship” or other user defined method

v. -p: Bonferroni correction, variants whose p-values above 0.05/number of tests will not be written to the file.

c. Input file:

i. genotype.hdf5

ii. phenotype.tsv

iii. genotype.kin.rescaled.IBS

d. Output file:

i. ./EMMAX_res/xxx.top

SKAT (Wu et al. 2010; Wu et al. 2011) was download as an R package (https://cran.r-project.org/web/packages/SKAT/index.html) and the p-values for regions were obtained by first computing the parameters and residuals for SKAT using following command line a).

a) >> obj<-SKAT_Null_Model(y ~ 1, out_type="D"), where y denotes phenotype matrix, out_type=”D” means the phenotype is dichotomous.

To perform the association studies between the SNPs set and the phenotype, we used the command line b)

b) >> res_p_value <- SKAT(x, obj)$p.value. Here, obj is generated by either a) or b) and x refers to genotype matrix for all SNPs in the SNPs set. “res_p_value” is the p-value for a tested SNP set associated with phenotype y. Please refer to the manual of SKAT for more details.

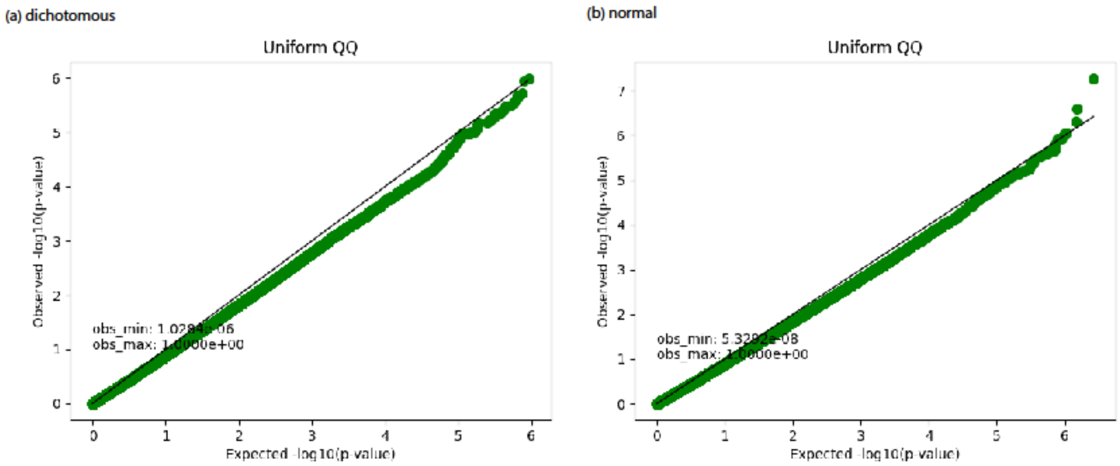


**Supplementary Figure S1.** Uniform QQ plot for simulated phenotype under null hypothesis. (a): Dichotomous phenotype (0 or 1); (b): Phenotype from normal distribution (mean zero and standard deviation 1).

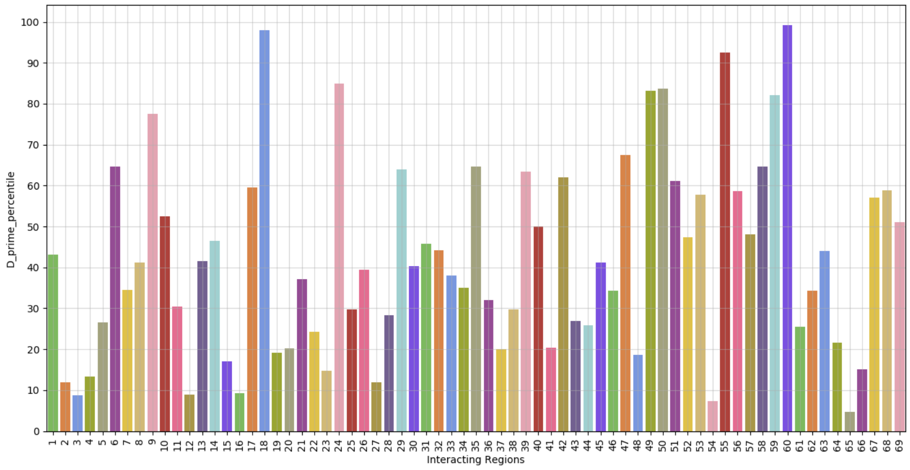


**Supplementary Figure S2.** D’ values for 69 interacting regions associated with both ASD and gene expressions in the brain tissues.
